# Molecular imaging reveals a high degree of cross-seeding of spontaneous metastases in a novel mouse model of synchronous bilateral breast cancer

**DOI:** 10.1101/2020.11.27.401786

**Authors:** Shirley Liu, Nivin N Nyström, John J Kelly, Amanda M Hamilton, Yanghao Fu, John A Ronald

## Abstract

Synchronous bilateral breast cancer (SBBC) patients present with cancer in both breasts at the time of diagnosis or within a short time interval. They show higher rates of metastasis and lower overall survival compared to women with unilateral breast cancer. However, the lack of a preclinical model has led to a dearth in knowledge regarding the patterns of SBBC metastasis. Here we established an SBBC model and used molecular imaging to visualize the development of spontaneous lung metastases arising from each primary tumor. We engineered human breast cancer cells to express either Akaluc or Antares2 for bioluminescence imaging (BLI), and tdTomato or zsGreen for *ex vivo* fluorescence microscopy. Both cell populations were implanted into contralateral mammary fat pads of mice (n=10) and BLI was performed weekly for up to day 29 (n=3), 38 (n=4), or 42 (n=3). Signal from both Antares2 and Akaluc was first detected in the lungs on day 28 and was present in 9 of 10 mice at endpoint. *Ex vivo* fluorescence microscopy of the lungs revealed that for mice sacrificed on day 38, a significant percentage of micrometastases were composed of cancer cells from both primary tumors (mean 37%; range 27% to 45%), while two mice sacrificed on day 42 showed percentages of 51% and 70%. These results reveal a high degree of metastatic cross-seeding which may contribute to faster metastatic growth and intratumoral heterogeneity. We posit our work will help understand treatment resistance and optimal planning of treatment for SBBC patients.

## INTRODUCTION

Synchronous bilateral breast cancer (SBBC) is defined as the detection of breast cancer in both breasts at the time of diagnosis or within a short interval of 3-12 months^1^. SBBC accounts for 2-11% of all breast cancer cases, and the estimated risk of SBBC development is 2-6 times higher in patients already diagnosed with breast cancer^2, 3^. Several clinical studies have found that compared to patients with unilateral breast cancer (UBC), SSBC patients have higher rates of distant metastasis and lower overall survival^2–7^. In 2015, Jobsen and colleagues reported that the aggressiveness of primary tumors in SBBC patients does not appear to differ from UBC patients, as measured by malignancy grading, mitotic activity index, hormone receptor status, and the presence of positive lymph nodes^2^. They suggested that the worse prognosis of SBBC is due to the combined effect of two tumors, resulting in a higher chance of metastasis. Moreover, when adjusting for disease characteristics such as tumor size, malignancy grade, histological subtype, nodal involvement, and estrogen receptor (ER)-status of both tumors, instead of only the worst tumor, SBBC does not show a higher mortality rate compared to UBC, supporting the idea that poorer outcomes are caused by the effect of two simultaneous tumors^7^. These studies provide important insight on why SBBC patients have worse prognoses; however, little is known about the spread and growth of metastatic cells from each primary tumor over time.

Animal models of UBC have existed for decades and have provided vital information on the steps, kinetics, and mechanisms of breast cancer metastasis, as well as the development and testing of antimetastatic treatments^8–12^. Similar animal models of SBBC are lacking. Thus, to complement the data from clinical SBBC studies, we developed a spontaneous metastases mouse model of SBBC where each mouse formed two primary breast cancer tumors in contralateral mammary fat pads. Cancer cells implanted into each mammary fat pad were pre-engineered to express a unique set of imaging reporter genes allowing us to distinctly visualize the metastatic fate of cancer cells from each tumor using non-invasive *in vivo* dual-bioluminescence imaging (BLI) and *ex vivo* fluorescence microscopy.

BLI relies on the reaction between luciferases and exogenously delivered substrates to produce light that is then detected by a charge-coupled device^13^. In this study, we used two highly sensitive BLI reporters Antares2 and Akaluc to improve our ability to monitor SBBC metastasis in deep tissues. Antares2 is an enhanced version of the original Antares reporter developed by Dr. Michael Lin’s group. Antares is a bioluminescence resonance energy transfer (BRET) reporter protein consisting of the BLI reporter NanoLuc fused to two copies of the orange fluorescent protein CyOFP1^14^. Reacting with furimazine, a synthetic analog of the Renilla luciferase substrate coelenterazine, NanoLuc emits blue light that can be absorbed by CyOFP1 and re-emitted as orange light, allowing better tissue penetration. Antares2 was developed by replacing NanoLuc with a mutant named teLuc^15^. Akaluc is a recently described luciferase that also promotes red-shifted light emission^16^. This BLI reporter was developed by Dr. Atsushi Miyawaki’s group through successive rounds of mutagenesis of Firefly luciferase to optimize pairing with the substrate AkaLumine hydrochloride^16^. Akaluc has been shown to be able to achieve detection of single cells at depth (lungs) in mice^16^. In addition to these BLI reporters, the fluorescence reporters zsGreen and tdTomato were co-expressed by either Antares2 or Akaluc-expressing cells respectively, to enable *ex vivo* microscopic cell characterization of SBBC metastases^17^.

Using this first SBBC model, we demonstrate that the vast majority of individual lung metastases are formed from cancer cells derived from both primary tumors rather than from one individual tumor, highlighting a high degree of cross-seeding. If present in SBBC patients, this high level of cross-seeding from distinct primary tumors may contribute to increased metastatic growth, tumor heterogeneity and treatment resistance. Future work using our model and potential variants may help further elucidate the biological mechanisms underlying the worse prognosis of SBBC patients.

## MATERIALS AND METHODS

### Lentiviral Construction and Production

Third generation lentiviral packaging and envelope-expression plasmids pMDLg/pRRE, pRSV-Rev, and pMD2.G were gifts from Didier Trono (Addgene plasmids #12251, #12253, and #12259, respectively)^18^. pUltra-Chili-Luc, a third-generation lentiviral transfer vector encoding tdTomato (tdT) and firefly luciferase (FLuc) separated by a P2A self-cleaving peptide sequence, was a gift from Malcolm Moore (Addgene plasmid #48688). All cloning was performed using In-Fusion HD Cloning (Takara Bio USA, Inc). As previously reported^19^, we built a pEF1α-tdT/FLuc2 transfer vector by replacing FLuc with FLuc2 and the Ubiquitin C promoter with a human elongation factor 1 alpha promoter (pEF1α) in the pUltra-Chili-Luc vector. To generate a pEF1α-tdT/Akaluc transfer plasmid, the Fluc2 sequence in pEF1α-tdT/FLuc2 was replaced with Akaluc obtained from the pcDNA3-Venus-Akaluc Vector (Cat. RDB15781, RIKEN BioResource Research Center). To generate a pEF1α-zsG/Antares2 transfer plasmid, the pEF1α-tdT/FLuc2 vector was modified to replace tdT with the fluorescence reporter zsGreen 1 (zsG) obtained from the pLVX-ZsGreen1-N1 Vector (Cat. 632565, Takara Bio USA, Inc.) and FLuc2 was replaced with the Antares2 sequence obtained from pcDNA3-Antares2 c-myc, a gift from Huiwang Ai (Addgene plasmid #100027)^15^. To produce pEF1α-zsG/Antares2 and pEF1α-tdT/Akaluc lentiviruses, the packaging, envelope and one of the transfer plasmids were co-transfected into human embryonic kidney (HEK 293T) cells using Lipofectamine 3000 (ThermoFisher Scientific) according to the manufacturer’s lentiviral production protocol. Lentivirus-containing supernatants were harvested 24- and 48-hours post transfection, filtered through a 0.45-μm filter, and stored at −80°C prior to use.

### Cell Culture and Lentiviral Transduction

Human triple negative breast cancer cells (MDA-MB-231) were obtained from a commercial supplier (American Type Culture Collection; ATCC) and cultured in DMEM supplemented with 10% fetal bovine serum at 37°C and 5% CO2. All cells were routinely verified as free of mycoplasma contamination using the MycoAlert mycoplasma detection kit (Lonza). Cells were transduced with pEF1α-zsG/Antares2 lentivirus overnight in the presence of 4-to 8-μg/mL polybrene. Transduced cells were washed, collected, and sorted for zsG expression using a FACSAria III fluorescence-activated cell sorter (BD Biosciences), generating Antares2-expressing cells with >95% purity. A second population of MDA-MB-231 cells were transduced with the pEF1α-tdT/Akaluc lentivirus and sorted for tdT expression to generate Akaluc-expressing cells with >95% purity.

### Furimazine and AkaLumine-HCl

Furimazine was purchased as NanoLuc substrate in the Nano-Glo® Luciferase Assay System (Promega) and was diluted 50x in media for *in vitro* experiments and 50x in PBS for *in vivo* experiments, as previously described^20^. AkaLumine-HCl (TokeOni; Sigma-Aldrich) was diluted in PBS to 5 mM for *in vivo* experiments. Further dilutions to 250 μM were made in media for *in vitro* experiments.

### *In Vitro* Studies

Fluorescence microscopy was performed on an Olympus IX50 Inverted System Microscope to visualize zsG in Antares2-expressing cells and tdT in Akaluc-expressing cells. To assess Antares2 and Akaluc activity and the correlation between cell number and BLI signal, increasing cell numbers from 1×10^4^ to 1.5×10^5^ cells were seeded in a 24-well plate. The next day, media was removed and replaced with 200 μL of furimazine diluted 50x in media for Antares2-expressing cells, or 200 μL of 250 μM AkaLumine-HCl in media, and images were acquired 5 minutes later.

To evaluate the substrate *in vitro* cross-reactivity of furimazine with Akaluc and AkaLumine-HCl with Antares2, two 24-well plates were plated with 5×10^4^ Akaluc-expressing cells, Antares2-expressing cells, naïve cells, and equivalent volume of media. The next day, media was removed and replaced with 200 μL of 250 μM AkaLumine-HCl in media in one plate, and 200 μL of furimazine diluted 50x in media in another plate. Images were acquired 5 minutes after incubation. After initial images were acquired, cells were washed with PBS, incubated with 1 mL of fresh media, and signal decay was measured for up to three hours. Cells were washed again every 24 hours followed by BLI until signal reached background levels. Results were obtained from three independent experiments with three replicates for each condition. The mean signal across replicates was determined for each independent experiment.

### *In Vivo* Studies

Animals were cared for in accordance with the standards of the Canadian Council on Animal Care, and under an approved protocol of the University of Western Ontario’s Council on Animal Care (2017-032). Six to eight-week-old female nu/nu mice were obtained from Charles River Laboratories (Willington, MA, USA).

We first evaluated the kinetics of Antares2 and Akaluc *in vivo* in nu/nu mice. Mice received orthotopic injections of either Antares2-expressing cells (5×10^5^; n=4) or Akaluc-expressing cells (5×10^5^; n=4) into the right fourth mammary fat pad (day 0). On day 5, all mice received an intravenous injection of 100 μL of furimazine (diluted 50x in PBS) and images were acquired every 60 seconds for 30 minutes, as well as at 1, 2, 3, and 24 hours. On day 6, all mice received intraperitoneal injections of 100 μL of 5 mM AkaLumine-HCl in PBS, and images were acquired every 60 seconds for 30 minutes, then at 1, 2, 7, 8, 11, 24, and 36 hours. BLI was performed on an IVIS Lumina XRMS In Vivo Imaging System (PerkinElmer). For all imaging, mice were anesthetized with 1-2% isofluorane using a nose cone attached to an activated carbon charcoal filter for passive scavenging. Regions of interest (ROIs) were manually drawn around tumor borders using LivingImage software to measure bioluminescent average radiance (p/s/cm^2^/sr).

To generate a SBBC model, eight-week-old female nod-scid-gamma (NSG) mice were obtained from an in-house colony (Dr. David Hess; Western University). Each mouse received orthotopic injections of 3×10^5^ Antares2-expressing cells into the right fourth mammary fat pad and 3×10^5^ Akaluc-expressing cells into the left fourth mammary fat pad (day 0; n=10). Antares2 BLI was performed weekly starting day 0 for up to 6 weeks. Images were acquired for up to 10 minutes. Akaluc BLI was performed on the same or on consecutive days after scans taken prior to Akalumine-HCl injection confirmed lack of Antares2 signal. Images were acquired for up to 30 minutes. Mice 1-3 were sacrificed on day 29 upon detection of both Antares2- and Akaluc-expressing lung metastasis and tissues were collected for histological analysis. The remaining seven mice were monitored for primary tumor growth and metastasis until endpoint, determined by the development of primary tumor necrosis (day 4-2 for mice 46 and day 38 for mice 7-10).

### Histology

At endpoint, mice were sacrificed by isoflurane overdose and perfused with 4% paraformaldehyde (PFA) via the left ventricle. Mammary fat pad tumors and lungs were removed and fixed in 4% PFA for an additional 48 hours, cryopreserved in ascending concentrations of sucrose (15 and 30% w/v) for 24 hours each. Tissues were then immersed in OCT medium (Sakura Finetek), frozen using dry ice, and sectioned at 14-μm thickness. Nuclei were stained with Hoechst and fluorescence microscopy of sections was performed on an EVOS FL Auto 2 Imaging System to evaluate zsG and tdT expression.

To analyze cell distribution of metastases from each primary tumor, individual micrometastases were identified in the lung. Micrometastases were identified as clusters of cells with a diameter larger than 200 μm, distinct from isolated tumor cells with a diameter less than 200 μm. The number of micrometastases composed of cells expressing zsG only, tdT only, or both zsG and tdT were manually counted in 5 fields of view per lung section for 3 lung sections for two mice sacrificed on day 42 and four mice sacrificed on day 38.

### Statistics

Linear regression analysis was performed to determine the goodness-of-fit coefficient (R^2^ value) for BLI signal versus number of Akaluc- or Antares2-expressing cells. Unpaired two-tailed t tests were performed using Graphpad Prism software (Version 8.1.2 for Mac OS X, GraphPad Software Inc., La Jolla California USA, www.graphpad.com) to compare BLI signal from Akaluc- and Antares2-expressing cells for *in vitro* and *in vivo* experiments. A *p*-value less than 0.05 was considered statistically significant. For imaging measurements over time, one-way ANOVA and Tukey’s multiple comparison post hoc test was performed.

## RESULTS

### Antares2 and Akaluc Do Not Exhibit Substrate Cross-reactivity

We first generated lentiviral vectors encoding either a pEF1α-zsG/Antares2 or a pEF1α-tdT/Akaluc cassette (Figure 1A). Human breast cancer cells were transduced with each lentivirus and sorted for zsG or tdT to obtain Antares2- and Akaluc-expressing cell populations with >95% purity (Figure 1B). Fluorescence microscopy also confirmed zsG and tdT expression in purified Antares2-expressing and Akaluc-expressing cell populations, respectively (Figure 1C). As expected, Antares2-expressing cells also exhibited some red fluorescence due to CyOFP1 in the fusion protein. To test for Antares2 and Akaluc functionality, increasing numbers of cells were seeded in a well plate and BLI signal was measured after co-incubation with the appropriate substrate (Figure 1D/E). A significant linear correlation was found between BLI signal and the number of both Antares2-expressing cells (Figure 1D; n=3, R^2^=0.9810, p<0.0001) and Akaluc-expressing cells (Figure 1E; n=3, R^2^=0.9944, p<0.0001).

**Figure 1.**
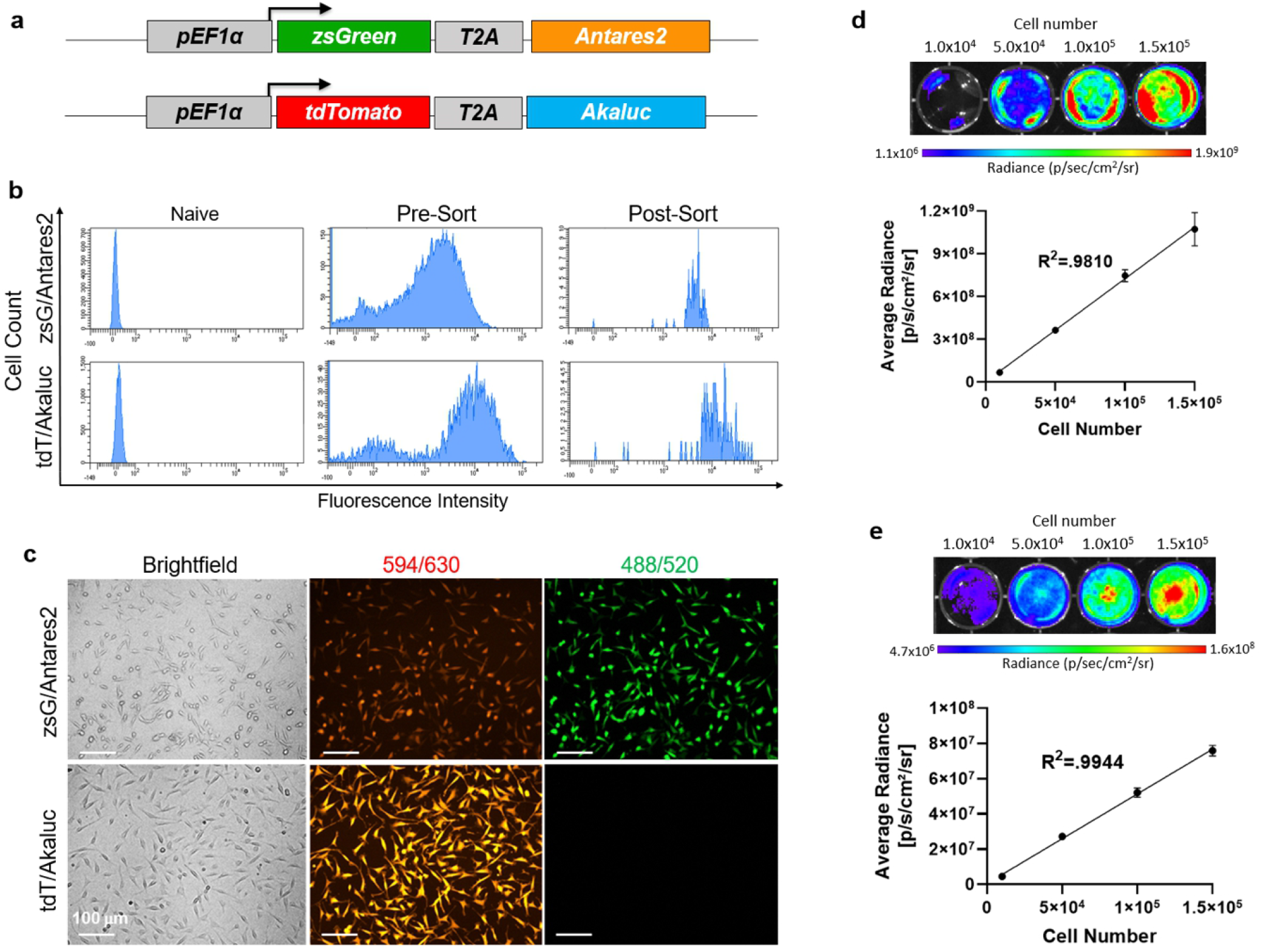
Lentiviral transduction of MDA-MB-231 cells to express fluorescent and bioluminescent reporter genes: (a) Reporter gene constructs for coexpression of zsGreen (zsG) and Antares2, and tdTomato (tdT) and Akaluc. (b) Histograms of control (naïve) and transduced MDA-MB-231 cells fluorescence-activated cell sorted for zsG or for tdT. (c) Fluorescence microscopy of sorted MDA-MB-231 cells engineered to express zsG/Antares2 or tdT/Akaluc, with excitation laser/emission filter wavelengths. (d) Bioluminescence imaging (BLI) signal vs number of Antares2-expressing cells after administration of furimazine. (e) BLI signal vs number of Akaluc-expressing cells after administration of AkaLumine-HCl. The data are presented as mean ± SD.

To validate the use of Antares2 and Akaluc reporters for dual-BLI of SBBC growth and metastasis, we first needed to confirm a lack of substrate cross-reactivity of furimazine with Akaluc and AkaLumine-HCl with Antares2. Akaluc-expressing cells, Antares2-expressing cells, naïve cells, and an equivalent volume of media were incubated with furimazine. Antares2-expressing cells showed significantly higher BLI signal (10^3^-fold) than Akaluc-expressing cells, naïve cells, and media (Figure S1A; n=3, p<0.05). Similarly, when treated with AkaLumine-HCl, Akaluc-expressing cells showed a significantly higher BLI signal (10^3^-fold) than Antares2-expressing cells, naïve cells, and media (Figure S1B; n=3, p<0.01). We next characterized the *in vitro* signal kinetics to determine the time for Antares2 and Akaluc signal to decay to background levels. Antares2-and Akaluc-expressing cells were treated with the appropriate substrate and imaged 5 minutes later. Cells were then washed with PBS and incubated in fresh media without substrate, and images were acquired over time with additional washes every 24 hours until negligible signal remained. Antares2-expressing cells showed negligible signal remaining by 24 hours after substrate treatment (Figure S1C). In contrast, Akaluc-expressing cells showed some signal remaining at 24 hours, but negligible signal by 48 hours (Figure S1D). This data suggests that Antares2 and Akaluc dual-BLI can be performed on consecutive days for *in vitro* experiments, with Akaluc BLI following Antares2 BLI.

We next evaluated the substrate cross-reactivity between Antares2 and Akaluc *in vivo*. Nude mice were implanted with either Antares2- or Akaluc-expressing cells into the mammary fat pad (n=4 per cell line). We allowed the tumors to grow and become palpable before conducting cross-reactivity tests. On day 5, furimazine was injected intravenously in all eight mice and BLI signal from mammary fat pad tumors were measured. On day 6, prescans confirmed the loss of Antares2 signal if previously present, and AkaLumine-HCl was injected intraperitoneally in all mice and BLI signal was measured. Antares2 tumors showed significantly higher signal (10^2^-fold) than Akaluc tumors when mice were injected with furimazine (Figure S2A; n=4, p<0.01). Similarly, Akaluc-expressing tumors showed significantly higher signal (10^4^-fold) than Antares2-expressing tumors after injection of AkaLumine-HCl (Figure S2B; n=4, p<0.01). These data validate the lack of substrate cross-reactivity between Antares2 and Akaluc both *in vivo* and support their use in dual-BLI. For *in vivo* kinetics studies, nude mice bearing Antares2 or Akaluc mammary fat pad tumors were injected with the appropriate substrate and imaged over time until negligible signal remained. Antares2 BLI signal peaked immediately and dropped to background levels by 3 hours after furimazine injection (Figure S2C). In contrast, Akaluc BLI signal peaked at 30 minutes and dropped to background levels by 36 hours after AkaLumine-HCl injection (Figure S2D). From these results, *in vivo* Antares2 and Akaluc dual-BLI could be performed on the same day or on consecutive days, with Akaluc BLI following Antares2 BLI.

### Dual-BLI of Contralateral Tumors Shows Spontaneous Metastasis of Both Antares2- and Akaluc-expressing Cells to the Lungs

To generate a SBBC mouse model, Antares2-expressing cells were implanted into the right mammary fat pad and Akaluc-expressing cells into the left mammary fat pad of ten NSG mice. We had six mice in our first cohort (mice 1-6) and four mice in a second cohort (mice 7-10). Dual-BLI was performed on the same or consecutive days weekly to track the growth of primary tumors and development of spontaneous metastases. Mice were imaged until endpoint as determined by the first presence of both Antares2 and Akaluc BLI signal in the lungs (day 29 for mice 1-3), or when evidence of ulceration of the primary tumors was present (day 38 for mice 7-10 and day 42 for mice 4-6).

On day 0, only the Antares2 tumor was visualized upon administration of furimazine, and on day 1 only the Akaluc tumor was visualized upon administration of AkaLumine-HCl (Figure 2A). Average radiance of BLI signal significantly increased over time for both Antares2 and Akaluc mammary fat pad tumors (Figure 2B; p<0.05).

**Figure 2.**
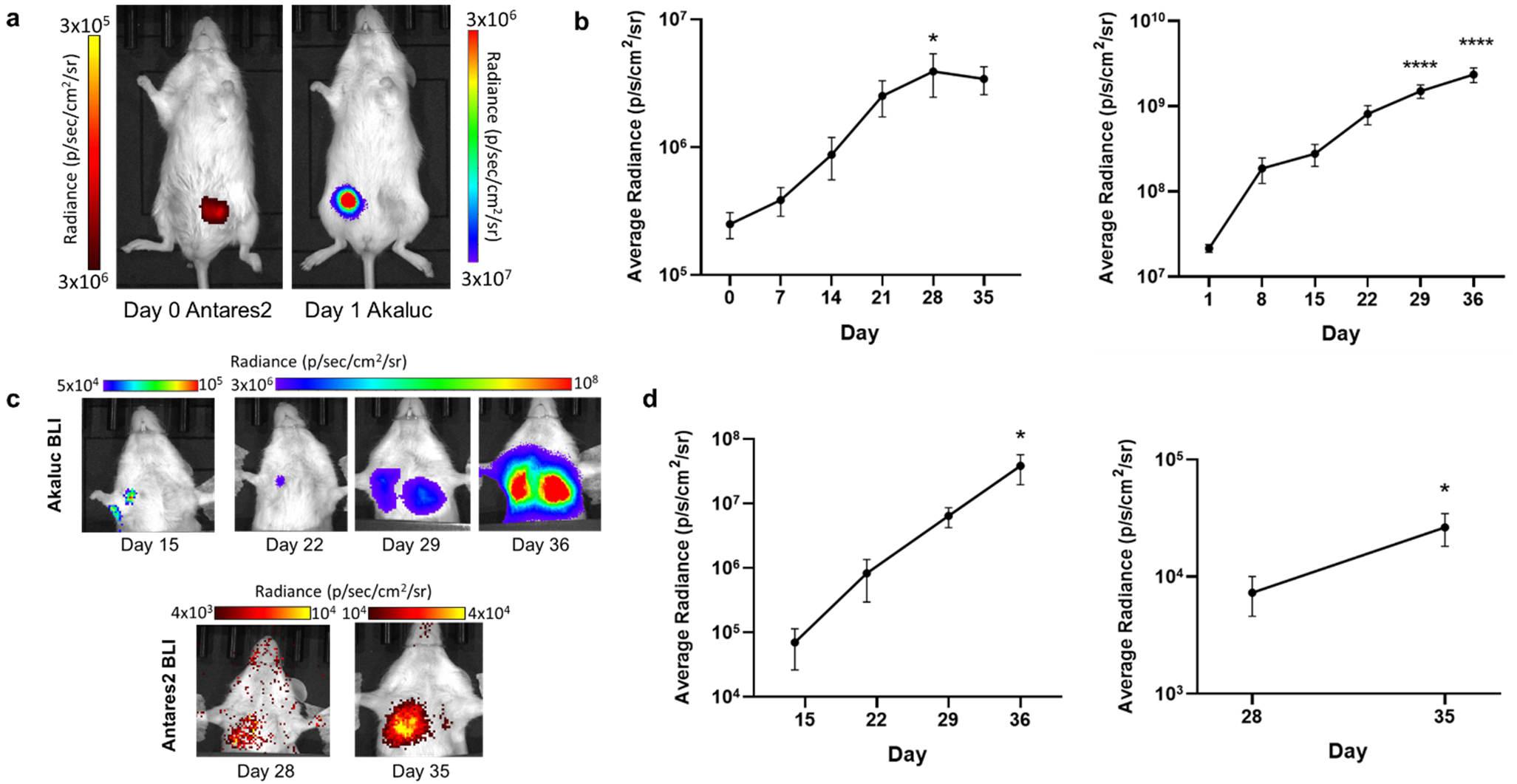
Dual-bioluminescence imaging (BLI) of Antares2 and Akaluc mammary fat pad tumors and lung metastasis in nod-scid-gamma mice: (a) Representative images of a mouse bearing contralateral mammary fat pad tumors imaged with Antares2 BLI on Day 0 and Akaluc BLI on Day 1. (b) Quantification of Antares2 (left) and Akaluc (right) mammary fat pad tumor BLI signal over time (n=10). (c) Representative images of Akaluc and Antares2 BLI of lung metastasis. (d) Quantification of Akaluc (left) and Antares2 (right) lung BLI over time (n=10). *p<0.05, ****p<0.0001 when compared to the initial data point. The data are presented as mean ± SEM.

Metastases were first detected in the lungs in six of the 10 mice on day 15 using Akaluc-BLI, with all mice showing Akaluc signal in the lungs by day 29 (Figure 2C). Antares2-BLI detected lung metastasis in four of the 10 mice by day 28, and in six of seven mice on day 35. By endpoint, nine of the 10 mice showed both Antares2 and Akaluc signal in the lungs. Average radiance significantly increased over time for both Antares2 and Akaluc BLI signal from the lungs (Figure 2D, p<0.05).

### A High Percentage of Micrometastases are Composed of Cancer Cells from both Primary Tumors

At endpoint, primary tumors and lungs were collected for analysis. Of the 10 mice, one mouse (mouse 4) was not preserved properly and thus excluded from further analysis. Akaluc primary tumors showed only tdT fluorescence, while Antares2 primary tumors showed only zsG fluorescence (Figure S3). We believe the absence of visible orange-red CyOFP1 fluorescence in Antares2-expressing cells is due to the perfusion fixation of the mice.

The lungs of the 3 mice sacrificed on day 29 showed both zsG- and tdT-expressing cancer cells, which were mostly found in unique loci and isolated from each other (Figure 3A/B). No micrometastases, defined as lesions with a diameter larger than 200 μm, had yet developed in these lungs, yet both Antares2 and Akaluc signal was clearly visible. For mice sacrificed on day 38 (mice 7 through 10), there were micrometastases present, with many composed of zsG-expressing cells, a moderate fraction composed of both zsG- and tdT-expressing cells, and a small fraction composed solely of tdT-expressing cells (Figure 3A/B, Figure S4). Finally, for the two mice sacrificed on day 42 (mice 5 and 6), many micrometastases were present in the lungs, with a large fraction qualitatively composed of both zsG- and tdT-expressing cells (Figure 3A/B).

**Figure 3.**
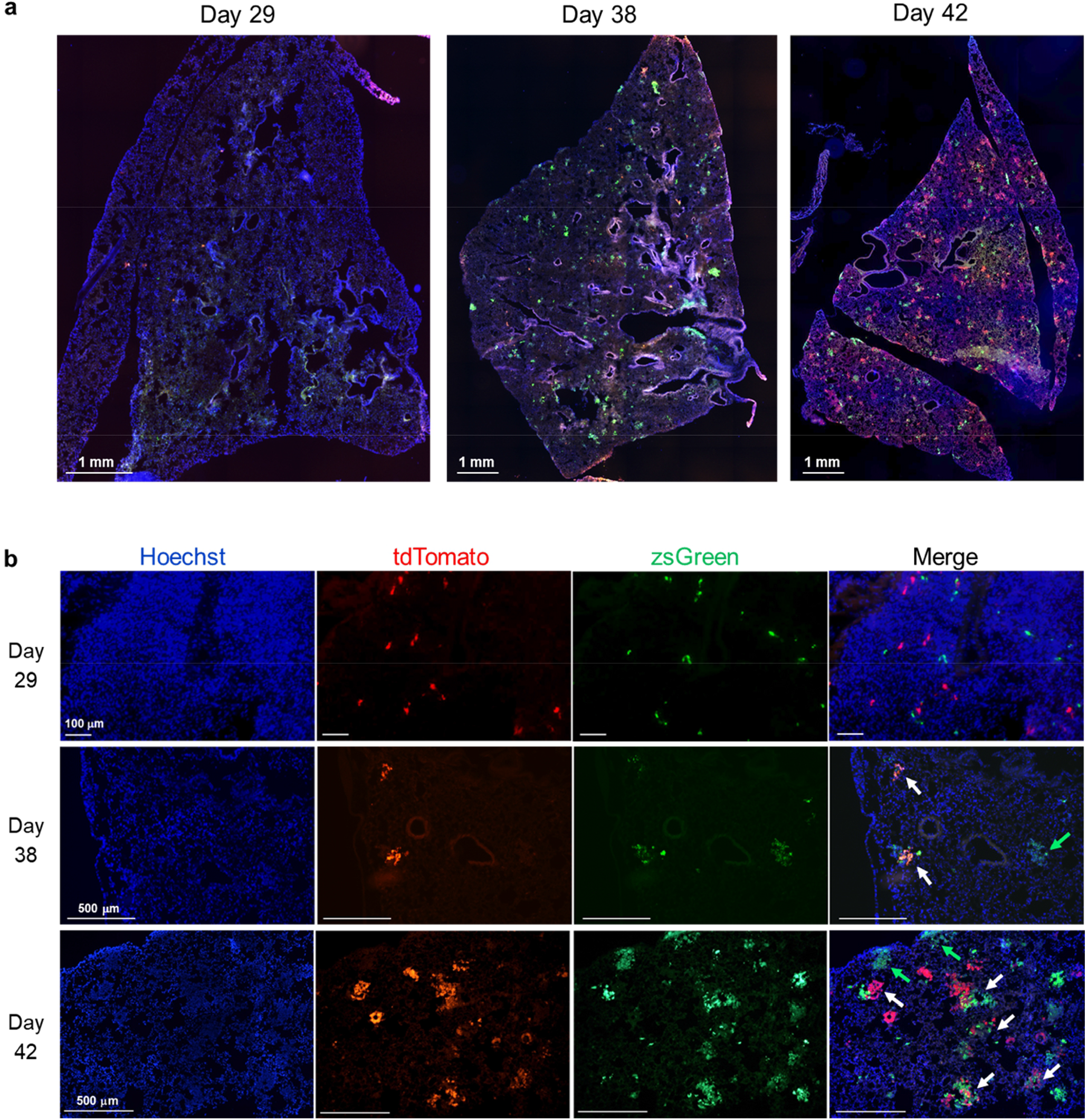
Fluorescence microscopy lungs of mice sacrificed on day 29, day 38, and day 42: (a) Representative whole lung sections of mice sacrificed at the three endpoints showing the presence of metastatic tdTomato (tdT) and zsGreen (zsG)-expressing cells. (b) Higher magnification images (10x) showing micrometastases composed of cells derived from both mammary fat pad tumors (white arrows) and micrometastases composed of only zsG-expressing cells (green arrows). No micrometastases composed of only tdT-expressing cells were identified in these fields of view.

To quantify the composition of micrometastases, the number of lung micrometastases in mice 5 through 10 composed of only tdT-expressing cells, only zsG-expressing cells, or both cell types were manually counted in five fields of view in each of three lung sections (Figure 4A). For mice 7-10 sacrificed on day 38, the mean percentage of micrometastases composed of both cell types was 37%, with percentages ranging from 27% to 45% (Figure 4B). For mice 5 and 6 sacrificed on day 42, 51% and 70% of micrometastases were composed of both cell types, respectively. When averaged across all six mice, there was a significantly higher percentage of micrometastases composed of both cell types compared to the percentage of micrometastases composed of only tdT-expressing cells (Figure 4C; p<0.001). We noticed that for mice 7-10, there was a large proportion of zsG-only micrometastases compared to tdT-only micrometastases. We then analyzed the primary tumor masses of the mice at endpoint, and found that while mice 5 and 6 had similar zsG/Antares2 and tdT/Akaluc tumor masses, for mice 7-10 the masses of the zsG/Antares2 tumors were significantly higher than the masses of the tdT/Akaluc tumors (Figure 4D/E; p<0.001), likely explaining the high percentage of zsG-only micrometastases in this particular cohort.

**Figure 4.**
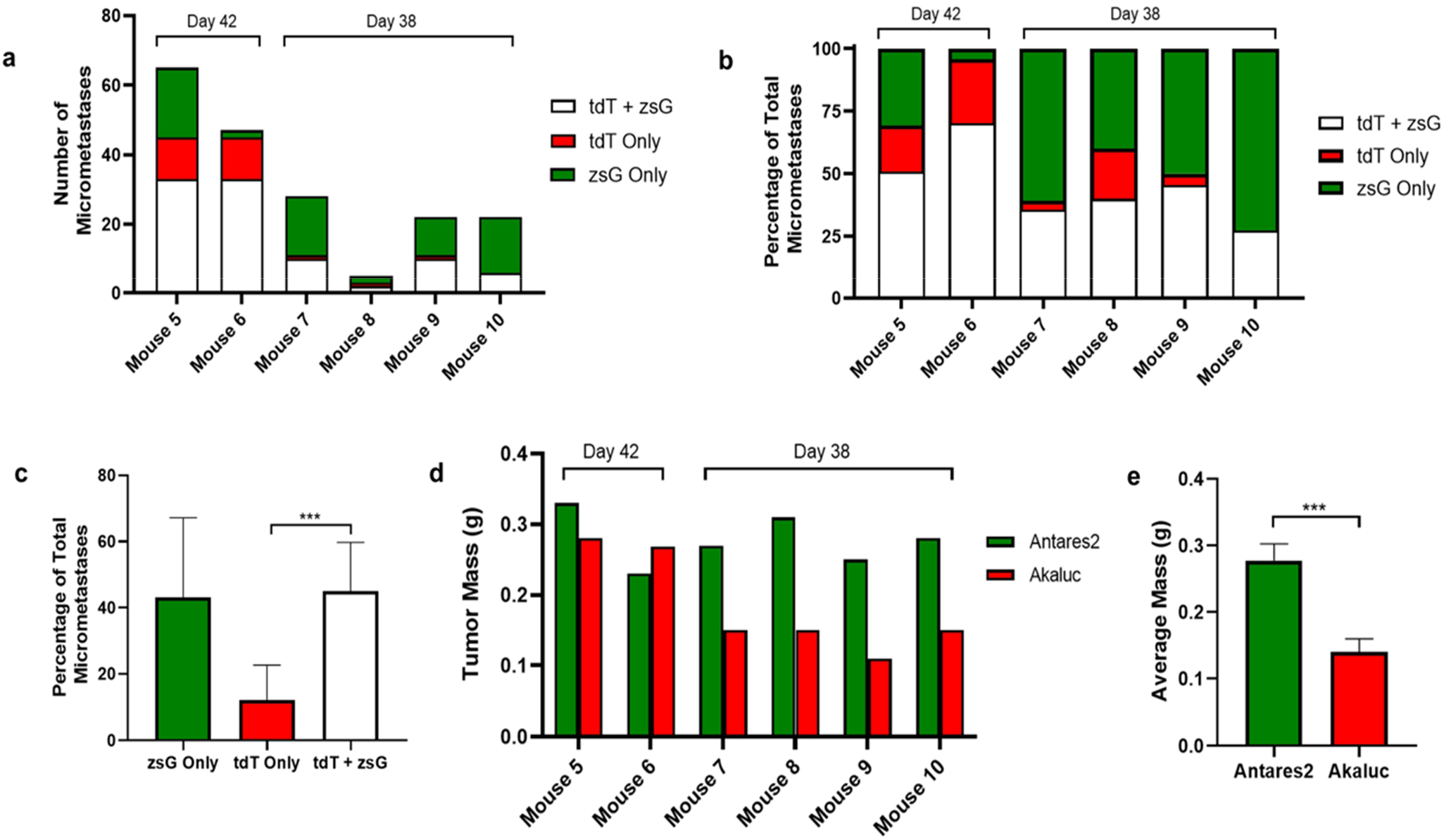
Cell composition analysis of lung micrometastases. The number of micrometastases (>200 μm diameter) composed of only zsGreen (zsG)-expressing cells, only tdTomato (tdT)-expressing cells, or both were counted in 5 fields of view at 3 lung sections for mice 5-6 (sacrificed on day 42) and mice 7-10 (sacrificed on day 38). Quantification of the types of micrometastases are expressed as total counts (a) or percentages of the total count (b). (c) Mean percentages of micrometastases types for mice 5-10 (n=6, ***p<0.001). (d) Mammary fat pad tumor masses for individual mice at endpoint. (e) Average mammary fat pad tumor masses for mice 7-10 (n=4, ***p<0.001). The data are presented as mean ± SD.

## DISCUSSION

SBBC is being presented to clinicians as an emerging concern, with increasing incidence due to improved screening technologies, prolonged life expectancy, and a growing awareness of the disease^1, 3^. Several retrospective clinical studies indicate that SBBC patients have higher rates of distant metastasis and higher mortality rates^2, 3^. Analysis of disease characteristics of SBBC and UBC have found no significant difference in their aggressiveness, leading to the hypothesis that the major explanation for worse prognosis of SBBC patients is an increased risk of metastasis as a result of having two cooperating bilateral tumors. However, due to the lack of longitudinal data characterizing the growth of SBBC metastases, little is known about the patterns of metastasis, undermining our ability to understand why SBBC patients are at increased risk of death. Characterizing the dynamics of SBBC metastasis will deepen our understanding of this disease and may eventually offer insights into mechanisms that may be targeted by anti-metastatic therapies, treatment resistance, and ways to better manage SBBC patients.

In this study we developed the first animal model to characterize metastatic SBBC. We genetically engineered human breast cancer cells to express either Antares2 or Akaluc for BLI, and zsG or tdT for fluorescence microscopy. We then implanted these cell populations into contralateral mammary fat pads of mice and used *in vivo* dual-BLI and *ex vivo* fluorescence microscopy to analyze the cell distribution of lung metastases arising from each mammary fat pad tumor. Signal from both Akaluc and Antares2-expressing cells was first visible in the lungs on day 28. By their respective endpoints, nine of 10 mice showed metastasis of both mammary fat pad tumors to the lungs. Analysis of the locations of these cells in mice sacrificed on day 42 showed that a majority of micrometastases in the lungs were composed of both zsG- and tdT-expressing cells, indicating metastasis of cells from both primary tumors to the same locations or the cross-seeding between zsG or tdT lung metastases once formed. We then performed this experiment again due to having only two mice sacrificed on day 42. In this second cohort of mice, due to the size of the primary tumours, these mice were sacrificed on day 38. Again, at endpoint we saw a significant portion of micrometastases composed of both zsG- and tdT-expressing cells, but compared to mice sacrificed on day 42, more micrometastases were composed of only zsG-expressing cells. We believe this to be due to faster-growing zsG/Antares2 mammary fat pad tumors in this particular cohort compared to the first cohort, as shown by a significantly higher average primary tumour mass at endpoint. When averaged across all six mice, we found that the percentage of micrometastases composed of both cell types was significantly higher than the percentage of micrometastases composed of only tdT-expressing cells.

Our results support previous findings that the worse prognosis of SBBC patients compared to UBC patients is likely not a result of more aggressive tumors, but due to the effects of two tumors combining to increase the progression of metastases^5^. However, our study provides new information on the patterns of metastasis, suggesting that two contralateral tumors may preferentially co-populate individual metastases in a process we call metastatic cross-seeding. This is similar to a phenomenon known as tumor self-seeding, in which circulating tumor cells (CTCs) that disseminate from a primary tumor travel through the circulation and seed back to the original tumor^21, 22^. This process has been hypothesized to be due to leaky microvasculature of the original tumor, providing a favourable environment to infiltrate into and populate. Similarly, CTCs have been shown to seed onto established metastatic sites possessing favourable survival conditions^23–25^. We have recently found that systemically administered CTCs can readily self-home to pre-established spontaneous metastases that have disseminated throughout the body, and used this to develop a novel cell-based theranostic^25^. Another possibility is that cells from metastases in the lungs disseminate again and seed onto other metastases in the organ, further contributing to mixed zsG- and tdT expression within the same tumor. In the future, it will be of value to study this phenomenon with more invasive imaging technologies, such as intravital fluorescence microscopy, in order to visualize this process in real time. These mechanisms supporting the metastatic cross-seeding phenomenon may be an explanation for the current study’s findings that cells are more likely to seed onto pre-established metastases, contributing to a high percentage of lung metastases composed of cells that originated from distinct primary tumors.

The cross-seeding of contralateral tumors may help explain the accelerated growth of SBBC metastasis and ultimately the poorer outcomes shown by retrospective clinical studies. Importantly, if the primary cancers are discordant in tumor biology including histopathologic classification, mutation status, hormone receptor status, and HER2 receptor status, metastatic cross-seeding may lead to increased intratumoral heterogeneity, adding another layer of complexity when choosing the optimal treatment plan. A study of the pathological profiles of 55 SBBC patients found estrogen receptor status, progesterone receptor status, and HER2 status discordance rates between the two primary tumours of 15%, 11%, and 8%, respectively^26^. With these factors being established biomarkers for expected treatment response and prognosis, hormone receptor discordance in SBBC patients has been associated with increased rate of metastasis and higher mortality at 5 years^27, 28^. A cohort study on the effect of adherence to the German national S3-guideline for adjuvant therapy on the survival outcome of bilateral breast cancer patients showed that patients who were treated guideline-adherent for only the primary index tumor suffered a significantly decreased recurrence-free survival compared to patients treated guideline-adherent for both tumors^29^. Although this study shows a need to maximize targeting of both lesions, the risk of adverse side effects of rigorous therapy plans must also be considered. The limited scientific evidence on the disease and a current lack of international guidelines on therapeutic management calls for a need to further investigate the properties of SBBC and the impact of various treatment plans on survival.

A limitation of our work is that we only characterized the cellular composition of metastases in the lungs where cells were located using *in vivo* BLI. Besides the lungs, SBBC patients also commonly show metastasis to the bone, liver, brain, and axillary lymph nodes^2, 5^. In our model, the growth of the primary tumors hindered our ability to image for a longer period of time. Surgically removing the primary tumors may allow us to prolong our study until metastasis in other organs can be detected. This has previously been demonstrated in a mouse model of breast cancer, where removal of the primary tumor extended the life span of mice but did not inhibit metastasis to distant organs^30^. Enhancing the sensitivity of our BLI reporters may also enable detection of other metastases. The sensitivity of Antares2 could be improved by pairing it with diphenylterazine (DTZ), a more optimal substrate^15^. Akaluc BLI could be improved by increasing the concentration of its substrate AkaLumine-HCl. By improving the sensitivity of Antares2 and Akaluc and covering the primary tumors and lungs to avoid saturation of the CCD camera, it may be possible to detect metastasis in other organs. Another limitation is that Antares2 shows some red fluorescence due to the presence of the orange fluorescent protein CyOFP1 in the fusion protein, which is visible in *in vitro* images. However, fluorescence microscopy of the Antares2 tumor did not show any perceptible red fluorescence, which we believe is due to the perfusion fixation of the mice, leading to reduced CyOFP1 fluorescence intensity. In addition, tdT-expressing cells do not show any green fluorescence, so all red fluorescence from lung sections is from tdT.

Future preclinical research on SBBC will aim to further characterize the mechanisms of metastasis and explore the effect of primary tumour discordance in hormone receptor and HER2 status. Models consisting of cells taken directly from both primary tumours in SBBC patients may allow us to better represent the tumor heterogeneity, although patient-derived xenograft (PDX) models are difficult to establish and the ability to establish PDX in both mammary fat pads may be even more difficult^31^. Future work in clinical research of SBBC could involve genome sequencing of tumors to track the lineages of metastatic cells. Mutational profiling has previously been performed to determine whether primary tumors in SBBC patients were of independent or clonal origin, which may have clinical implications^32^. Similarly, gene sequencing techniques could also be applied to elicit the origin of metastatic tumors and determine their heterogeneity.

### Conclusions

In this work we established the first animal model of SBBC and evaluated the metastatic cell distribution of contralateral mammary fat pad tumors using *in vivo* dual-BLI and *ex vivo* fluorescence microscopy. We found that the majority micrometastases in the lungs are composed of cells from both primary tumors, suggesting a high degree of metastatic cross-seeding which we posit may contribute to intratumoral heterogeneity and treatment resistance. Our work deepens our understanding of the mechanisms underlying poor outcomes of SBBC patients and may offer insight into optimal management of SBBC.

## Conflict of Interest

The authors declare no conflicts of interest.

## Funding

This research was funded by a Canadian Institutes of Health Research (CIHR) Project Grant held by J.A.R., grant number 377071. S.L. was supported by an Undergraduate Student Research Award from the Natural Sciences and Engineering Research Council of Canada.

## Notes

### Competing Interest Statement

The authors have declared no competing interest.

