## Supplementary Material for "Molecular imaging reveals a high degree of cross-seeding of spontaneous metastases in a novel mouse model of synchronous bilateral breast cancer"

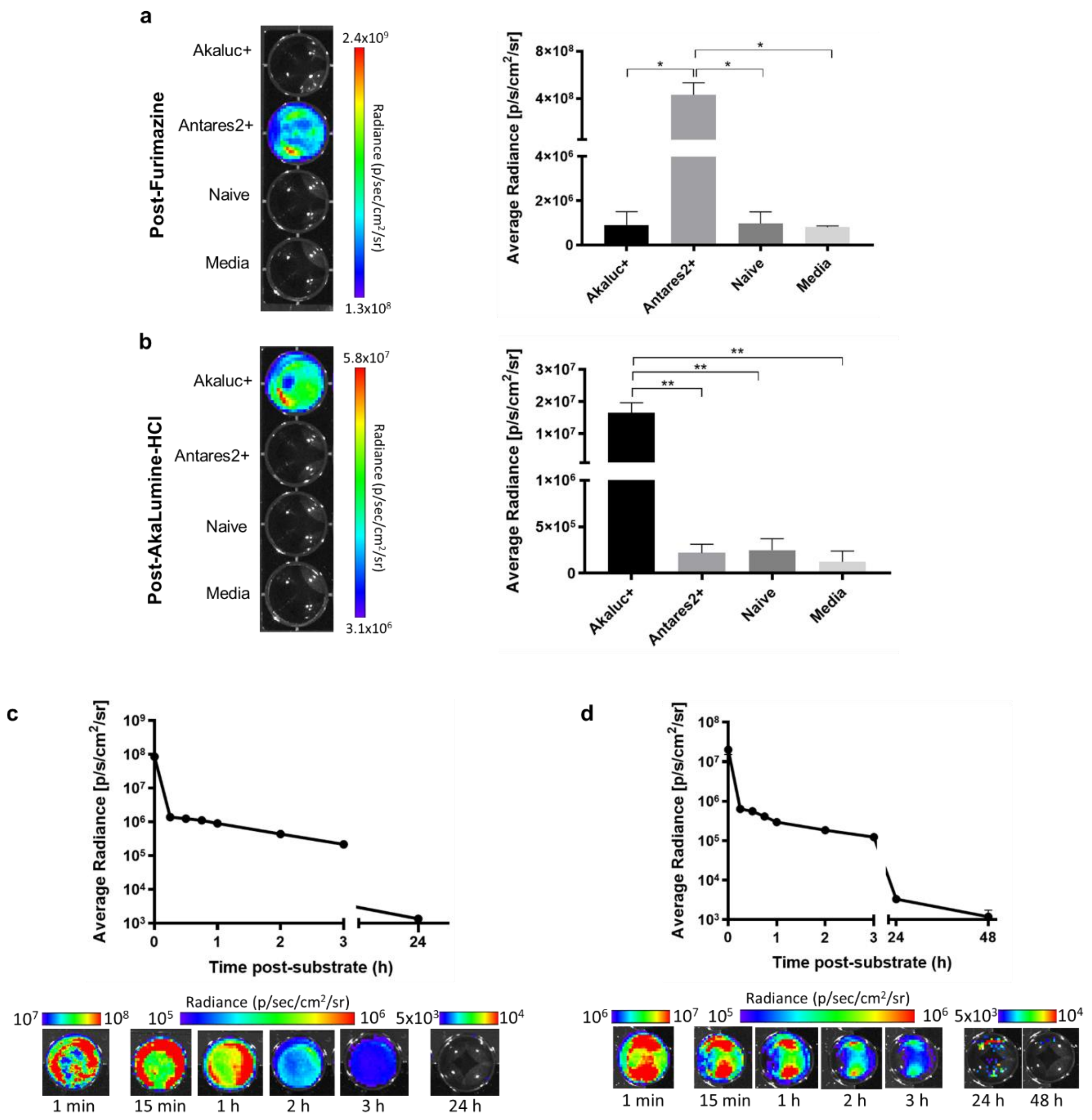


**Figure S1.** *In vitro* cross-reactivity and kinetics of Antares2 and Akaluc. Cells were treated with the indicated substrate at 0 hrs, then washed with PBS and incubated in media for subsequent images. Cells were washed with PBS every 24 hours until negligible signal remained: (a) Representative well plate and quantification of bioluminescence imaging (BLI) signal of Akaluc-expressing cells, Antares2-expressing cells, naïve cells, and an equivalent volume of media after administration of furimazine (n=3, *$p<0.05$). (b) Representative well plate and quantification of BLI signal of Akaluc-expressing cells, Antares2-expressing cells, naïve cells, and an equivalent volume of media after administration of AkaLumine-HCl (n=3, **$p<0.01$). (c) BLI signal of Antares2-expressing cells over time after administration with furimazine (n=3). (d) BLI signal of Akaluc-expressing cells over time after administration with AkaLumine-HCl (n=3). The data are presented as mean ± SEM. Error bars for some data points are smaller than the corresponding symbols.

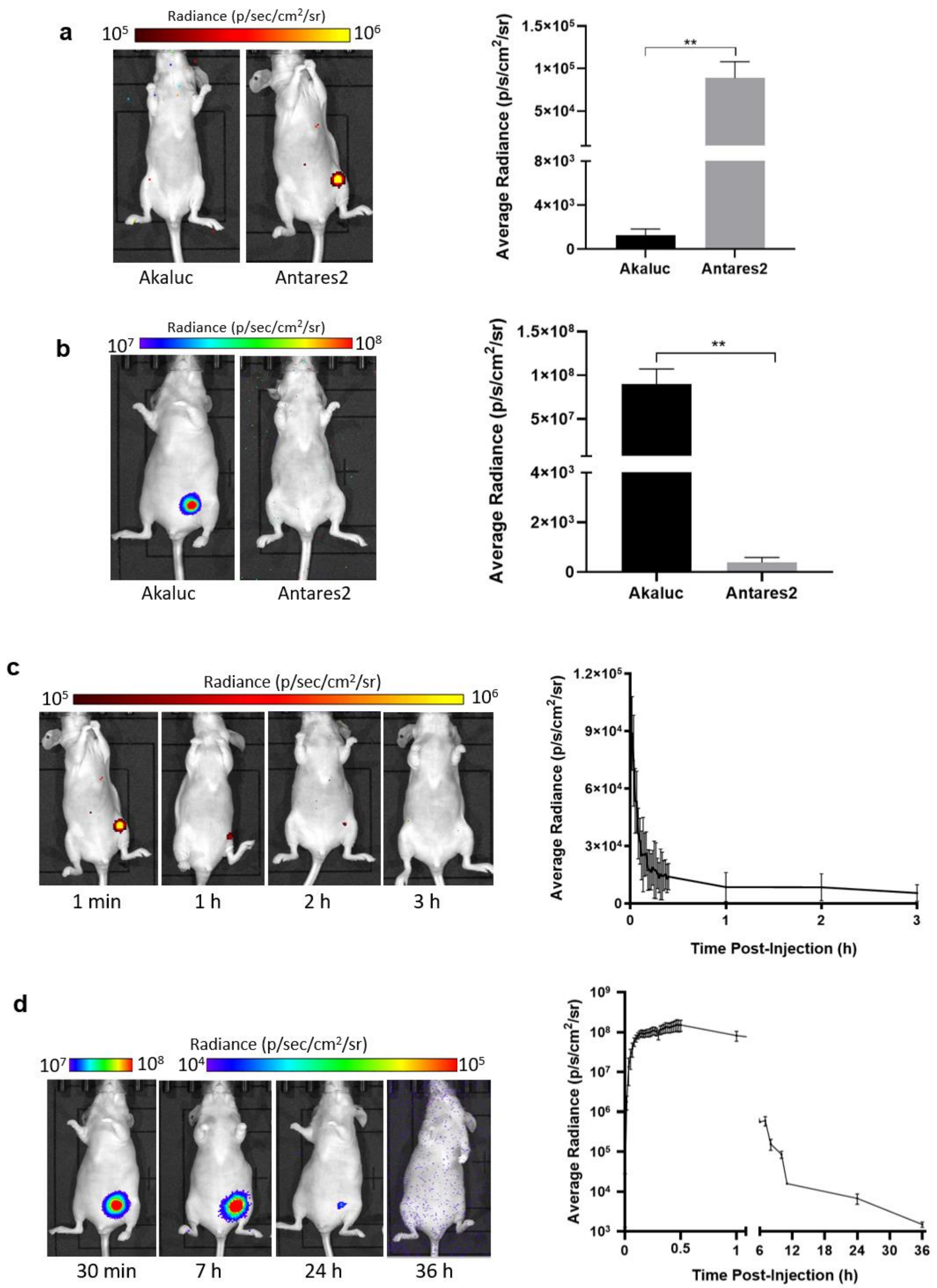


**Figure S2.** *In vivo* cross-reactivity and kinetics of Antares2 and Akaluc: (a) Representative bioluminescence imaging (BLI) images of a nude mouse bearing an Antares2 mammary fat pad tumor and injected intravenously with furimazine (n=4, **p<0.01). Images were acquired immediately and over time until negligible signal remained. (b) Representative images of a nude mouse bearing an Akaluc tumor and injected intraperitoneally with AkaLumine-HCl (n=4, **p<0.01). (c) Representative images and quantification of Antares2 BLI signal decay over time. (d) Representative images and quantification of Akaluc BLI signal decay over time. The data are presented as mean ± SEM.

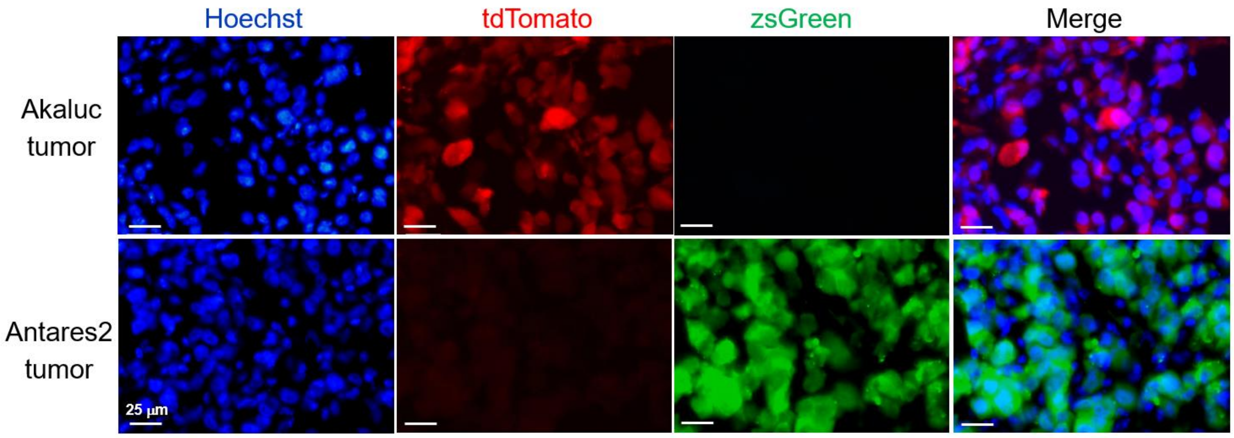


**Figure S3.** Fluorescence microscopy images of Akaluc and Antares2 mammary fat pad tumors of mice sacrificed on day 29.

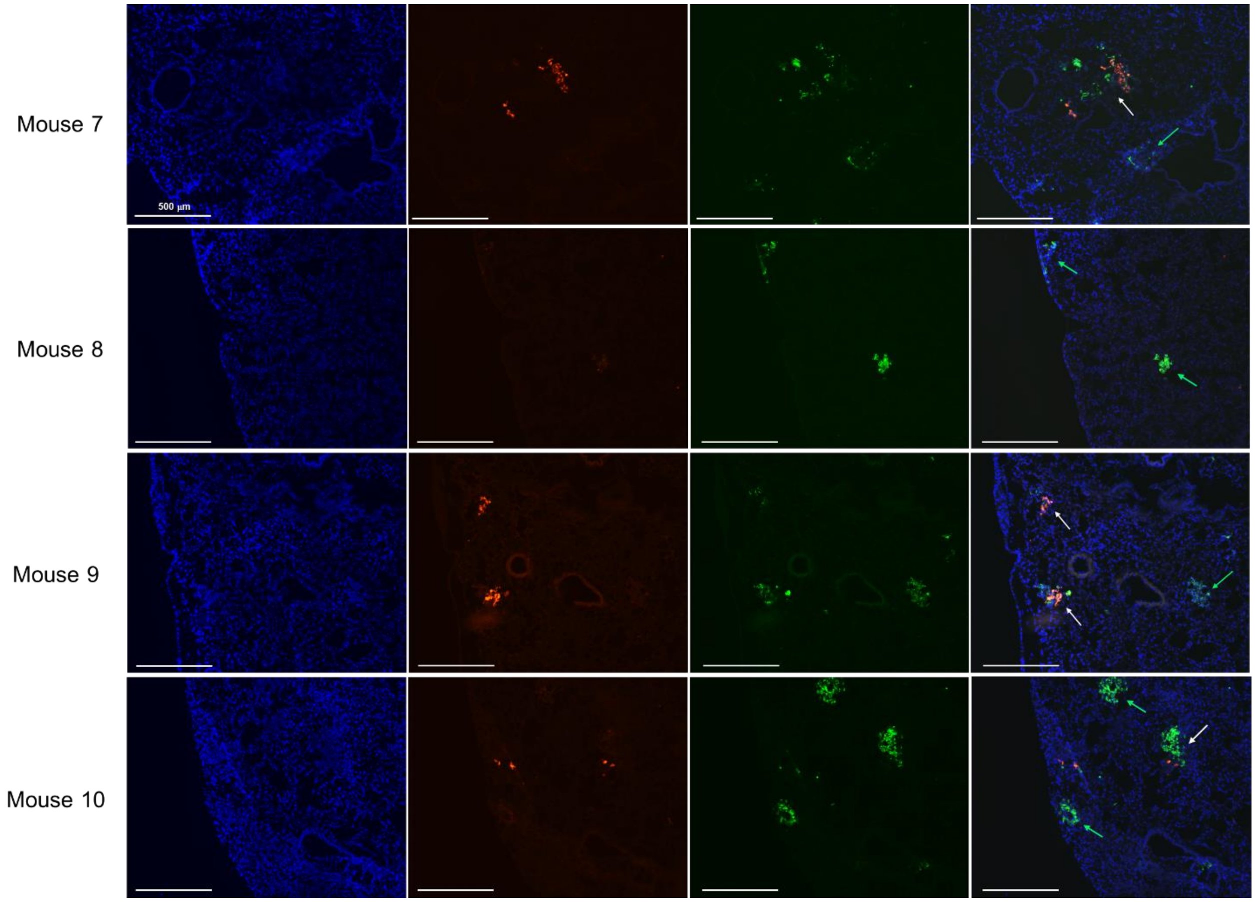


**Figure S4.** Fluorescence microscopy images of the lungs of mice sacrificed on day 38. Micrometastases (>200 μm diameter) composed of only zsG-expressing cells and both zsG- and tdT-expressing cells are indicated by green and white arrows, respectively. No micrometastases composed of only tdT-expressing cells were identified in these fields of view.
